# 2-Mercaptoethanol/DMSO workflow enables highly reproducible quantitative proteomics

**DOI:** 10.64898/2025.12.19.694883

**Authors:** Arisa Suto, Yoshihiro Ishikawa, Toshihide Matsumoto, Makoto Itakura, Yoshio Kodera, Takashi Matsui

## Abstract

Proteomics provides a systematic and high-throughput approach to comprehensively characterize protein networks, enabling insights into cellular functions and disease mechanisms. Carbamidomethylation using iodoacetamide (IAA), a common method for cysteine alkylation, is known to cause nonspecific modifications that increase MS spectral complexity and reduce quantitative accuracy. Here, we extended our 2-mercaptoethanol (2-ME) adduction method, enhanced by dimethyl sulfoxide (DMSO), from single-protein applications to a proteome-wide workflow. Mouse liver proteomes were processed using either 2-ME/DMSO or conventional IAA treatment, followed by LC-MS/MS analysis. The optimized 2-ME treatment increased the number of cysteine-modified peptides by 1.6–1.9-fold. Although total protein identifications were comparable, 77% of proteins exhibited improved sequence coverage. Quantitative reproducibility was also enhanced, with peptides quantified coefficient of variation (CV) ≤ 20% increasing from 61.5% (IAA) to 86.1% (2-ME), and proteins from 80.6% to 93.5%. Application of the workflow to an ovarian clear cell carcinoma reliably detected cisplatin-induced alterations. The 2-ME/DMSO workflow offers a simple and highly reproducible proteomics strategy for accurate quantitative proteomics.

## Introduction

Many biological processes in cells are orchestrated by proteins, and a comprehensive understanding of this complex network is essential for elucidating cellular functions and disease mechanisms. Proteomics provides a systematic and high-throughput approach to characterize the entire proteome of a biological sample, enabling an in-depth exploration of protein expression, modification, and interaction dynamics (1, 2). Therefore, this approach enables the identification of disease-associated pathways and clinically relevant biomarkers by providing quantitative insights into proteins whose abundances exhibit significant alterations under specific biological or pathological conditions (1, 2). In a typical proteomics workflow, disulfide bonds are cleaved using reducing agents such as dithiothreitol and tris-(2-carboxyethyl)-phosphine (TCEP) to facilitate the protein digestion(3). After reduction, thiol groups are protected by alkylating agents such as iodoacetamide (IAA). Subsequently, the reduced proteins are digested proteolytically using trypsin, and the resulting peptides are analyzed by liquid chromatography–mass spectrometry (LC-MS) to identify their sequence and measure their abundances (4). However, such alkylating agents are highly reactive with limited specificity, leading to offsite modifications at the N-terminus and various amino acids, particularly lysine, in addition to cysteine (5, 6). In particular, the reduction and alkylation steps influence the efficiency of enzymatic digestion and thereby lead to differences in the number of proteins identified depending on the combination of reducing and alkylating agents (7, 8). Consequently, the extent of side reactions caused by alkylating agents also varies substantially. Thus, these non-specific offsite modifications complicate MS spectra, lowering peptide identification rate, quantification accuracy, and sequence coverage. To address these limitations, we developed a cysteine-specific modification method intended for a purified single protein (9). This approach was inspired by unexpected observations that the reducing agent 2-mercaptoethanol (2-ME) modifies cysteine residues during in-gel digestion and protein crystallization (10, 11), and enhanced this reaction using dimethyl sulfoxide (DMSO), which is known to promote disulfide bond formation between cysteine-containing peptides(12). We previously developed a 2-ME/DMSO workflow that specifically modifies cysteine residues, thereby minimizing offsite modifications and improving both peptide identification and quantitative accuracy (9). Mechanistically, DMSO acts as an oxidizing agent and facilitates thiol oxidation and disulfide exchange, leading to the formation of reactive intermediates that readily undergoes the disulfide exchange reaction with 2-ME (12–14). Consequently, 2-ME preferentially forms mixed disulfides between cysteine residue and 2-ME, effectively suppressing unintended reactions at other nucleophilic sites.

Therefore, we extended the 2-ME adduction method to proteome-wide scale and examined whether this approach improves peptide identification and quantification reproducibility compared to conventional IAA alkylation. In this study, we optimized reaction conditions of proteome-wide 2-ME adduction and evaluated its performance using data-independent acquisition (DIA). We further applied the workflow to a biological sample, cisplatin-treated ovarian clear cell carcinoma (OCCC) model, as a use case.

## Materials and Methods

### Protein digestion of mouse liver

Male C57BL/6 mice were purchased from CLEA Japan, Inc. (Japan). All animal procedures were conducted in accordance with the guidelines of the National Institutes of Health and approved by the Animal Experimentation and Ethics Committee of Kitasato University School of Medicine(15). Proteins were extracted from the sliced mouse liver with a phase-transfer surfactant (PTS)(16), as previously described (17). The protein concentration of the homogenate was adjusted to 10 μg/μL based on absorbance at 280 nm measured using a NanoDrop spectrophotometer (Thermo Fisher Scientific, USA) in 50 mM Tris-HCl (pH 8.0).

For the 2-ME adduction, 20 μg of extracted protein was incubated with 20% DMSO (anhydrous) and 0-400 mM (0–2.8(v/v)%) 2-ME at 50°C for 30 min to promote thiol oxidation and disulfide exchange leading to 2-ME adduction.

For the carbamidomethylation, 20 μg of extracted protein was incubated with 22 mM Bond-Breaker TCEP solution (Thermo Fisher Scientific) at 50°C for 30 min to cleave the disulfide bond.

The disulfide bond cleaved protein was alkylated by 30 mM IAA at room temperature for 30 min in the dark for the carbamidomethylation at the cysteine residue, and then the reaction was quenched using 36 mM L-cysteine.

The 2-ME adducted or carbamidomethylated sample was digested using 200 ng trypsin and 200 ng lysylendopeptidase at 37°C overnight.

### Protein digestion of OVISE cell

The OVISE cells, derived from OCCC, were obtained from the National Institute of Biomedical Innovation (Japan). The cells were maintained in Eagle’s minimum essential medium or RPMI 1640 supplemented with 10% bovine calf serum. For anti-cancer drug stimulation, the cells were treated with 20 µM cis-diamminedichloroplatinum (II) (CDDP; Nippon Kayaku Co., Ltd., Japan) under 5% CO_2_ at 37°C, and the cell pellets were collected after 24 h according to the previously described(18). Proteins were extracted from the cell pellets with PTS (19) and the protein concentration was adjusted to 0.5 μg/µL by adding 50 mM Tris-HCl (pH 8.0) to a final volume of 20 µL.

For 2-ME adduction, 10 μg of the extracted protein was incubated with 20% DMSO and 300 mM 2-ME at 50°C for 30 min, followed by digestion with 100 ng trypsin and 100 ng lysylendopeptidase at 37°C overnight.

For the carbamidomethylation, 10 μg of extracted protein was incubated with 22 mM Bond-Breaker TCEP solution at 50°C for 30 min followed by alkylation using 30 mM IAA at room temperature for 30 min in the dark. After quenching the reaction mixture using 36 mM L-cysteine, the sample was digested with 100 ng trypsin and 100 ng lysylendopeptidase at 37°C overnight.

### Preparation for LC-MS analysis

The digested sample was mixed with 1.5 volumes of 1.7% trifluoroacetic acid (TFA), and the supernatant was collected after centrifugation at 19,000 × g at 4°C for 15 min. The supernatant was then loaded onto a MonoSpin C18 column (GL Sciences, Japan) for desalting, and the resulting peptides were eluted with 50% acetonitrile (ACN) containing 0.1% TFA, followed by freeze-dried.

### LC-MS/MS Data Acquisition

The LC-MS/MS analysis was performed using an Orbitrap Exploris 240 mass spectrometer (Thermo Fisher Scientific) equipped with a Vanquish Neo UHPLC system and the InSpIon system (AMR, USA) (20). The freeze-dried sample was resuspended in 3% ACN and 0.1% formic acid (FA), and digested peptides were injected directly onto an analytical column (C18, particle diameter 3 µm, 75 µm × 125 mm; Nikkyo Technos, Japan).

For Data Dependent Acquisition (DDA) analysis, the digested peptides were separated with a gradient of mobile phase A (0.1% FA) and mobile phase B (0.1% FA in 80% ACN), as follows: 0-0.1 min, 0-5.6%B; 0.1–20.1 min, 5.6–33.8%B; 20.1–26.1 min, 33.8–67.5%B; 26.1–27.1 min 67.5– 95%B, 27.1-30 min 95% B at a flow rate of 300 nl/min. The spray voltage was 2 kV and an ion-transfer tube temperature of 280°C. MS1 spectra were collected over the scan range 350–1200 *m*/*z* at 120,000 resolution to hit an automatic gain control (AGC) target of 3 × 10^6^ and a maximum injection time of 100 ms. The isolation width was set at 2 *m*/*z*, and the 20 most intense ions with charge states of 2^+^ to 4^+^ that exceeded an intensity of 2 × 10^3^ were fragmented by collision-induced dissociation with a normalized collision energy of 30%. MS2 spectra were acquired on the Orbitrap mass analyzer with a mass resolution of 15,000 to set an AGC target value was set at 5×10^5^ and a maximum injection time of Auto. The dynamic exclusion time was set to 15 s.

For DIA analysis, the digested peptides were separated with a 60 min gradient as follows: 0-5 min 0-5.6%B; 5–47 min, 5.6–33.8%B; 47–57 min, 33.8–62%B; 57–58 min 62–95%B, 58-60 min 95%B at a flow rate of 300 nl/min. MS1 spectra were obtained in the range of 495–775 *m*/*z* at 17,500 resolutions to set an AGC target of 3 × 10^6^ ions and a maximum injection time of 20 ms. MS2 spectra were collected at >200 *m*/*z* at 30,000 resolutions to set an AGC target of 3 × 10^6^ ions, a maximum injection time of 100 ms, and collision energy of 26%. The isolation width for MS2 was 9 *m*/*z*, and window patterns of 500–770 *m*/*z* were used.

### Peptide identification and quantification

DDA-MS data were analyzed using FragPipe (v.22.0, https://fragpipe.nesvilab.org/) with the MSFragger search engine (v.4.1)(21, 22), Philosopher (v.5.1.1)(23) and IonQuant (v.1.10.27)(24). Database searching was implemented against the UniProt protein sequence database (mouse, proteome ID: UP000000589, 21755 entries). Cysteine carbamidomethylation (+57.02, IAA), 2-mercaptoethanol adduction (+76.00, 2-ME), and methionine oxidation (+15.99) were set as variable modifications. No modification was set as fixed. The following parameters were applied: specific strict trypsin (trypsin/P), precursor mass tolerance of 10 ppm and fragment mass tolerance of 0.02 Da with mass calibration and parameter optimization enabled, a maximum of 2 missed cleavages, and a minimal peptide length of 7 amino acids. All other parameters were left at their default settings. The peptide identification was filtered at a false discovery rate (FDR) <1%.

DIA-MS data were analyzed using DIA-NN (version: 2.1.0, https://github.com/vdemichev/DiaNN) (25). First, an *in silico* spectral library was generated from the UniProt protein sequence database (Mouse, proteome ID: UP000000589, 21755 entries; Human, proteome ID: UP000005640, 20,644 entries) using DIA-NN. The Parameters for the spectral library generation were as follows: digestion enzyme, trypsin; missed cleavages, 2; peptide length range, 7–45 amino acids; precursor charge range, 2–4; precursor *m*/*z* range, 495–775; and fragment ion *m*/*z* range, 200–1800. The following options were enabled: “FASTA digest for library-free search/library generation;” “deep learning-based spectra, RTs, and IMs prediction;” “N-term M excision;” and “Ox (M)”. For variable modifications, 2-ME adduction on cysteine (UniMod:303, +75.99 Da) and carbamidomethylation on cysteine (UniMod:4, +57.02 Da) were applied for mouse data, whereas only the 2-ME adduction (UniMod:303) was applied for human data. For the DIA-NN search, the following parameters were applied: mass accuracy and MS1 accuracy were both set to 10 ppm; match-between-run was enabled for OCCC and disenabled for Mouse liver; protein inference was based on genes; neural network classifiers were used in cross-validated mode; quantification was performed using the Legacy (direct) quantification mode (Mouse liver) or the high-precision QuantUMS strategy (OCCC); and cross-run normalization was set to “RT-dependent”. The precursor and protein FDR was 1%, default setting.

### Experimental Design and Statistical Rationale

For the mouse liver experiment, a single homogenized liver extract was prepared and divided into six aliquots (three for IAA and three for 2-ME), each processed independently to generate three technical replicates per condition. For the OCCC cisplatin experiment, cells were cultured in separate wells and processed separately, and MS datasets of OCCC model treated with or without CDDP were performed with three biological replicates in each experimental condition.

Data preprocessing, statistical analyses, and visualization were performed in Python (v3.12) using Google Colab. NumPy (v2.0.2) and SciPy (v1.16.2) were used for data processing, statistical testing, and hierarchical clustering, while scikit-learn (v1.6.1) was applied for additional clustering analyses. Protein intensities were log_2_-transformed before quantitative analysis.

For the quantitative analyses in the mouse liver samples, peptides and proteins were retained only when they were detected in all three replicates within each condition, ensuring that only consistently detected analytes were compared between IAA and 2-ME.

For the OCCC cisplatin model, proteins were included in the quantitative analysis if they were detected in at all three replicates within at least one experimental group. Missing values in the other group were imputed using random numbers drawn from a normal distribution (width, 0.3; downshift, 1.8), reflecting values below the detection limit.

### Data analysis

Figures–including volcano plots and heatmaps–were generated using matplotlib (v3.10.0), seaborn (v0.13.2) and biopython (1.86). Fold change between conditions was calculated based on the ratio of log_2_-transformed protein intensities for comparative analysis, and Welch’s *t*-test was performed for differential significance, visualizing the results by volcano plots. Proteins with |log_2_(fold change)| ≥1 and Welch’s *t*-test *p* value < 0.01 (–log_10_(*p* value) > 2.0) were considered significant and classified as upregulated or downregulated proteins, respectively. Coefficient of variation (CV) value was calculated from the dataset within quantified peptides or proteins. Pathway enrichment and gene set enrichment analyses (GSEA) were conducted using the GSEApy (v1.1.10; https://github.com/zqfang/GSEApy) (26) based on the KEGG 2021 Human database. Each group was analyzed with the Enrichr module of GSEApy to identify significantly enriched KEGG pathways. For each pathway, enrichment score, *p*-value, and combined score were used to assess significance, and the top 15 enriched pathways were visualized as bubble plots. The protein–protein interaction (PPI) analysis was performed using the STRING database (ver12.0, https://string-db.org/). The full STRING network was used, in which edges represent both functional and physical associations, with a minimum interaction confidence score of 0.40 (medium confidence). The MCL clustering was applied to identify functional modules within the PPI network.

### Western Blot

Total cellular proteins were isolated using RIPA buffer (Fujifilm Wako Pure Chemical Corporation, Japan) supplemented with HALT^®^ protease and phosphatase inhibitor cocktail (Thermo Fisher Scientific). Aliquots of the proteins were resolved by SDS-PAGE, transferred to PVDF membranes, and probed with primary antibodies coupled to the ECL detection system (Cytiva, USA), as described previously (27). The primary antibodies included HNF-1β (BD Biosciences, USA), Cyclin D1 (DAKO, Denmark), p53 (Leica, USA) at dilutions ranging from 1:1000–1:2000.

## Results

### Optimization of Reduction and Modification Conditions with 2-ME in the DDA analysis of Mouse liver

The reaction condition for 2-ME adduction intended for the proteome-wide sample was optimized by varying the concentration of 2-ME ranging from 0 to 400 mM, using DDA-MS with mouse liver. 2-ME treatment markedly increased the number of peptide identifications and improved protein sequence coverage compared with IAA treatment (Fig1). 2-ME-adducted peptides were detected in all samples treated with 2-ME, not only in previously reported purified samples (9) but also in protein mixtures such as mouse liver. The number of 2-ME-adducted peptides increased in a concentration-dependent manner (Fig1.A). Approximately 1,800 2-ME-adducted peptides were identified at 300 mM 2-ME, representing 1.9 times increase compared with IAA treatment. The number of peptides without cysteine residues also increased in a concentration-dependent manner, and at 2-ME concentrations above 100 mM, more peptides were identified compared to IAA (Fig. 1A). Assessment of the total number of unique peptides at each missed-cleavage level demonstrated that the 2-ME treated samples consistently yielded more peptide identifications than the IAA treated samples across all digestion states, including fully cleaved peptides as well as peptides with one or two missed cleavages (Fig. 1B). Although peptide identifications increased, the total number of identified proteins was only slightly higher in the 2-ME treatment than in the IAA treatment (Fig. 1C). Notably, 77% of proteins showed higher sequence coverage with 2-ME than IAA (Fig. 1D). Peptide characteristics–including peptide length, *m*/*z*, molecular mass, GRAVY score, and pI–showed similar trends between peptides identified from 2-ME and IAA treatments (Supplemental Fig. S1), indicating that 2-ME does not result in a distinct peptide identification profile. In contrast to the physicochemical property, 2-ME treatment enabled the identification of a greater number of cysteine-modified peptides, leading to an increase in the frequency of cysteine residues (Supplemental Fig. S1F). When focusing exclusively on peptides uniquely identified by either IAA- or 2-ME-treatment, the distributions of peptide length, *m*/*z*, and molecular mass in both IAA- and 2-ME-tretments were similar trends as seen in those of all identified peptides; however, peptides identified from 2-ME-treatment tended to be longer length and exhibited higher *m*/*z* and molecular mass compared with those identified from the IAA treatment (Supplemental Fig. S2). Although GRAVY scores showed no substantial differences from the overall trend, peptides identified from IAA treatment notably displayed a marked decrease the median of pI. A bias of amino acid frequency only revealed in the IAA treatment, increasing probabilities of aspartic acid and glutamic acid residues which contribute lower pI (Supplemental Fig. S2E–F). The difference of physicochemical property suggests that 2-ME treatment performs minimal preparation-induced bias within cysteine-modified and unmodified peptides.

**Figure 1.**
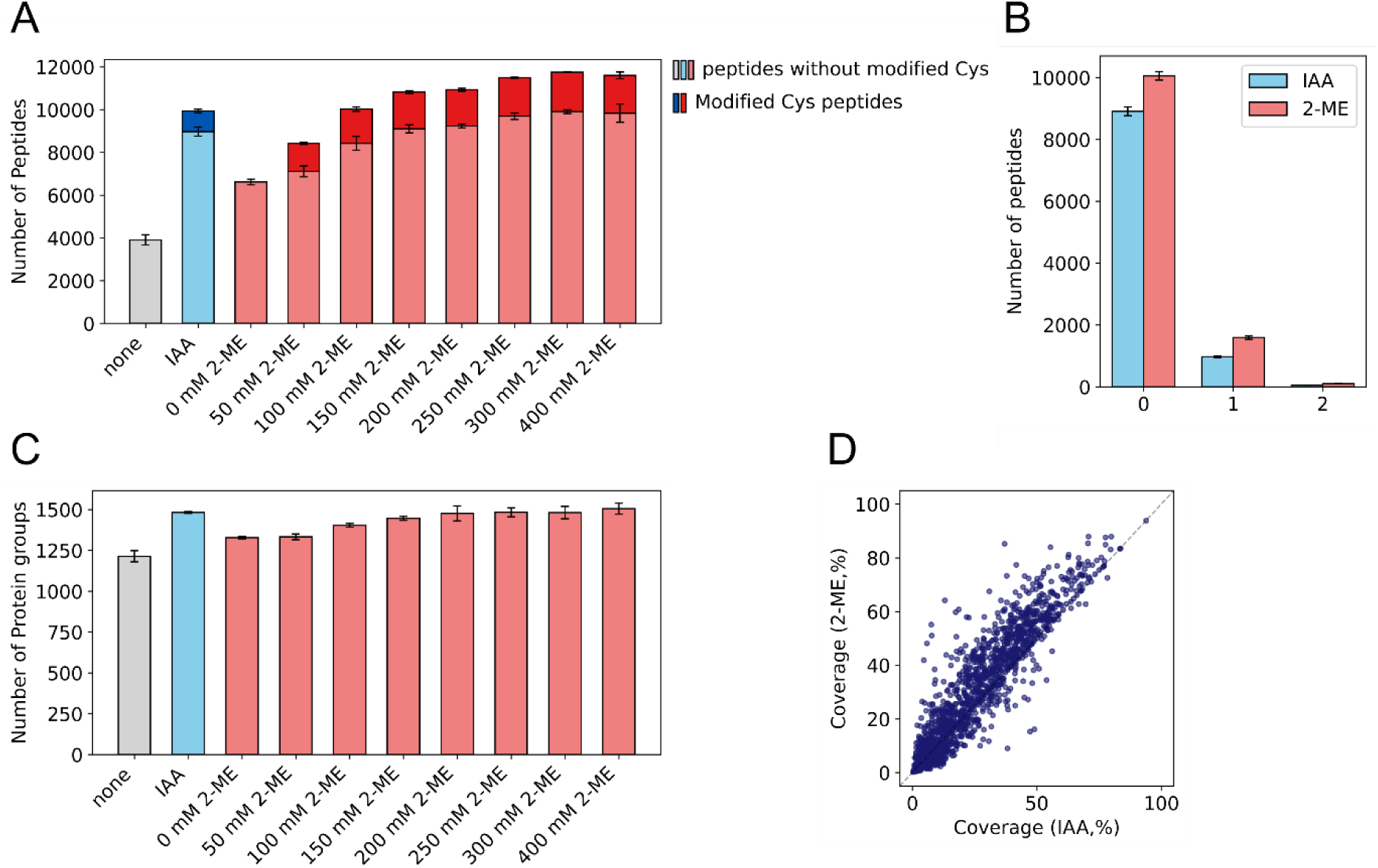
DDA analysis of mouse liver. A. Number of identified peptides in untreated samples (none) and in samples treated with IAA or various concentrations of 2-ME. Untreated, IAA-treated, and 2-ME-treated samples are shown in gray, blue, and red bars. Light- and dark-colored bars indicate peptides without or with peptides containing modified cysteine residue. B. Number of peptides at each missed-cleavage level (0-2) for IAA- and 2-ME-treated samples. C. Number of identified protein groups under the same conditions described in panel A. The values in panel A – C were presented as mean ± standard deviation. D. Scatter plot comparing protein sequence coverage between IAA and 300 mM 2-ME treated samples. Sequence coverage was calculated using the combined data from the three technical replicates within each condition.

### Evaluation of IAA and Optimized 2-ME Conditions in DIA analysis of Mouse Liver

To evaluate quantitative performance, we compared DIA analysis of samples processed with 300 mM 2-ME or 30 mM IAA. Although the total number of protein identifications did not differ between the two treatments, 2-ME treatment yielded a greater number and higher intensities of peptides, markedly cysteine-modified peptides, than IAA treatment (Figs. 2 and 3A–C). Moreover, 2-ME treatment improved quantitative reproducibility, as evidenced by a markedly increased proportion of peptides with CVs below 10–20% (Fig 3D–E).

**Figure 2.**
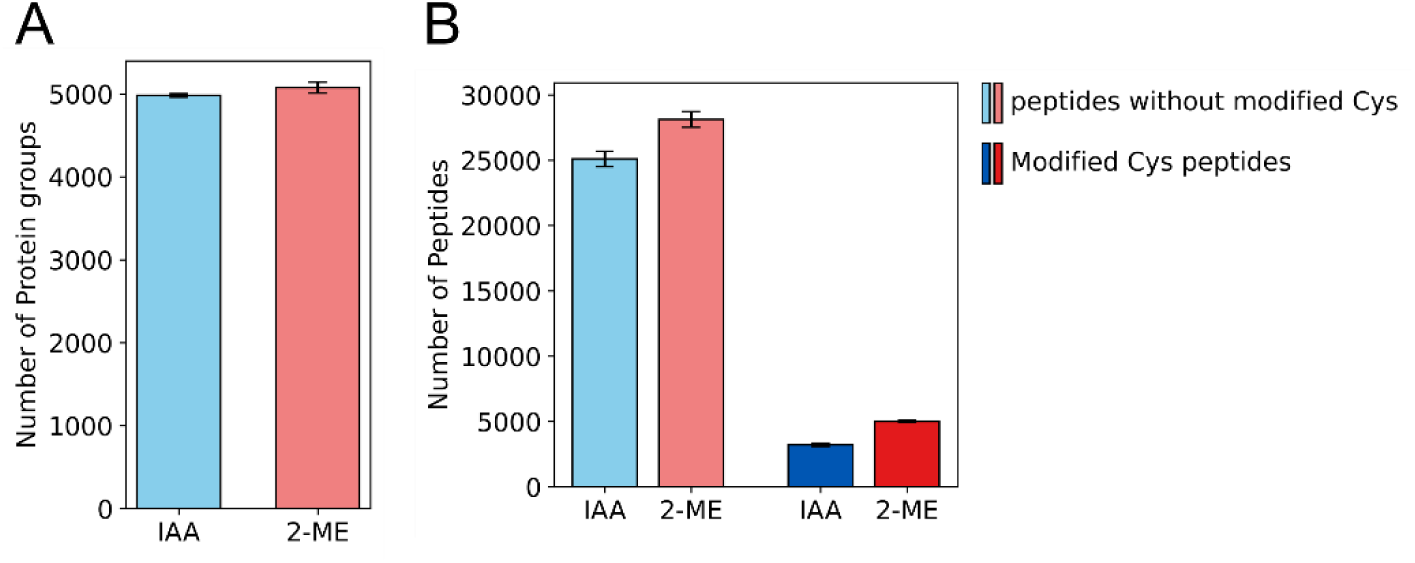
Number of identified proteins and peptides in mouse liver by DIA analysis. A. Number of identified protein groups for IAA- and 2-ME-treated samples. IAA- and 2-ME-treated samples are shown in blue and red bars. B. Number of identified peptides. Light- and dark-colored bars indicate peptides without or with peptides containing modified cysteine residues. All values were presented as mean ± standard deviation.

**Figure 3.**
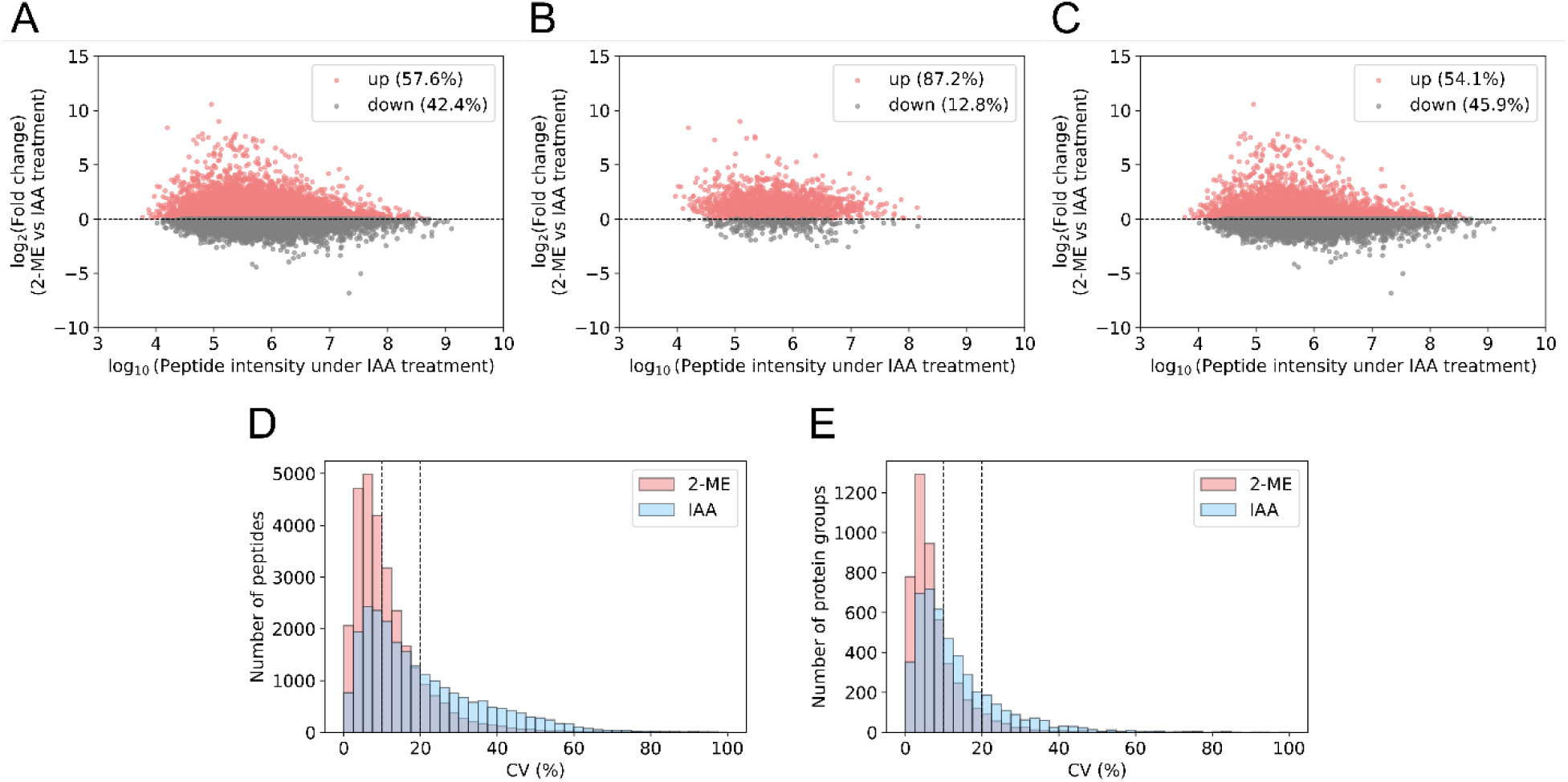
Peptide intensities and CV distribution by DIA analysis. A–C. Scatter plot comparing of log_10_ peptide intensities under IAA treatment with the log_2_ fold change of peptide intensities in 2-ME relative to IAA. A, B, and C show the whole peptides, cysteine-modified peptides, and peptides without modified cysteine residues, respectively. Peptides exhibiting increased intensities in 2-ME are highlighted in red. D–E. Comparative distributions of (D) peptides’ and (E) proteins’ CVs between IAA and 2-ME treated samples. Only peptides quantified in all three replicates are shown, with intensities averaged across replicates. CV values at 10% and 20% depicted by dotted lines, respectively.

The average number of identified proteins across three independent experiments was 4,980 for IAA-treated samples and 5,075 for 2-ME-treated samples, showing no substantial difference in identification numbers, consistent with the DDA analysis (Fig. 2A). Comparison of the identified proteins revealed that, among a total of 5,523 proteins, 5,114 proteins (92.6%) were detected in both IAA- and 2-ME-treated samples, indicating no significant difference in protein identification (Supplemental Fig. S3).

In contrast to the similarity observed at the protein level, the number of identified peptides differed markedly between treatments. Approximately 3,214 cysteine-modified peptides were detected in IAA-treated samples, whereas approximately 5,000 peptides, which is about 1.6-fold higher, were identified in 2-ME-treated samples (Fig. 2B). Furthermore, consistent with previous reports (9), the number of identified peptides without cysteine residues was also higher in 2-ME-treated samples, representing a 1.1-fold increase compared with approximately 25,091 peptides detected in IAA-treated samples (Fig. 2B).

Notably, peptides that showed low intensities under IAA treatment exhibited a marked increase following 2-ME adduction, including many peptides that increased by more than 2.5 log_2_ fold (∼5.6-fold) (Fig. 3A–C). This enhancement was particularly prominent for peptides near the detection limit. By improving signal intensity, 2-ME treatment directly contributed to the substantial improvement in quantitative reproducibility (Fig. 3D, Table 1). As a result, among cysteine-modified peptides, the proportion of peptides with CV ≤ 20% increased from 50.9% in the IAA-treated samples to 87.9% in the 2-ME treated samples, demonstrating a substantial improvement in reproducibility (Supplemental Fig. S4). Furthermore, considering all peptides, including those without cysteine residues, 61.5% of peptides in the IAA-treated samples were within CV 20%, whereas 86.1% of peptides in the 2-ME-treated samples met this criterion. In addition, in the 2-ME-treated samples, 15,945 peptides, corresponding to 56.3% of the total, fell within a CV of 10%, exceeding the 14,208 peptides that reached a CV of 20% in the IAA-treated samples (Figs. 3D and Supplemental Fig. S4). These results indicated that peptides showing a CV of 20% in IAA can be evaluated with a CV of 10% in 2-ME-treated samples, reflecting markedly higher reproducibility. Notably, at the protein level, the number of identified proteins with CVs below 20% increased by approximately 12.9% in the 2-ME-treated samples compared with IAA treatment, resulting in 93.5% of all identified proteins achieving CV ≤ 20% (Fig. 3E, Table 2). These results suggest that 2-ME treatment enables more accurate analyses in various comparative studies.

**Table 1.** CV of quantified peptides under IAA and 2-ME.

| Peptide type | IAA |  |  | 2-ME |  |  |
| --- | --- | --- | --- | --- | --- | --- |
|  | All | Modified Cys | Without Cys | All | Modified Cys | Without Cys |
| CV ≤ 10% | 7,489<br>(32.4%) | 490<br>(18.5%) | 6,999<br>(34.2%) | 15,945<br>(56.3%) | 2,492<br>(57.3%) | 13,453<br>(56.1%) |
| CV ≤ 20% | 14,208<br>(61.5%) | 1,351<br>(50.9%) | 12,857<br>(62.8%) | 24,384<br>(86.1%) | 3,823<br>(87.9%) | 20,561<br>(85.8%) |
| CV > 20% | 8,913<br>(38.5%) | 1,304<br>(49.1%) | 7,609<br>(37.2%) | 3,932<br>(13.2%) | 524<br>(12.1%) | 3,408<br>(14.2%) |

**Table 2.** CV of quantified proteins under IAA and 2-ME.

|  | IAA | 2-ME |
| --- | --- | --- |
| CV ≤ 10% | 2,381 (51.5%) | 3,581 (75.2%) |
| CV ≤ 20% | 3,724 (80.6%) | 4,453 (93.5%) |
| CV > 20% | 899 (19.4%) | 311 (6.5%) |

### Proteomic Analysis of cisplatin response in ovarian clear cell carcinoma

Because 2-ME treatment improved the peptide intensities, especially in the lower intensity region, and then enabled reproducible quantification with nearly all identified proteins exhibiting CV ≤ 20% (Fig. 3E and Table 2), we next examined whether this workflow is also effective for drug-response proteomics, where substantial protein loss often makes comprehensive quantification difficult. Therefore, we evaluated CDDP-induced proteomic alterations in OCCC cells.

A total of 7,571 proteins were identified under both CDDP-treated and untreated conditions, and notably, 97.2% (7,360 proteins) were successfully quantified. Among them, 81 proteins were upregulated and 467 proteins were downregulated following CDDP treatment (Fig. 4A–B). Western blotting validated proteomic changes, including upregulation of p53 (TP53) and Cyclin D1 (CCND1) and downregulation of HNF-1β (HNF1B) (Fig. 4C–F). Pathway enrichment and PPI analyses showed activation of mismatch repair (EXO1, SSBP1) and p53 signaling (CDKN2A, TP53), consistent with the well-known DNA-damage response caused by CDDP, which induces intra- and interstrand DNA cross-links that block replication and transcription (28) (Fig. 4G, Supplemental Fig. S5A). In contrast to the upregulated pathways, numerous ribosomal proteins were markedly downregulated, suggesting impaired protein synthesis and suppressed cell proliferation in response to CDDP treatment (Fig. 4H, Supplemental Fig. S5B).

**Figure 4.**
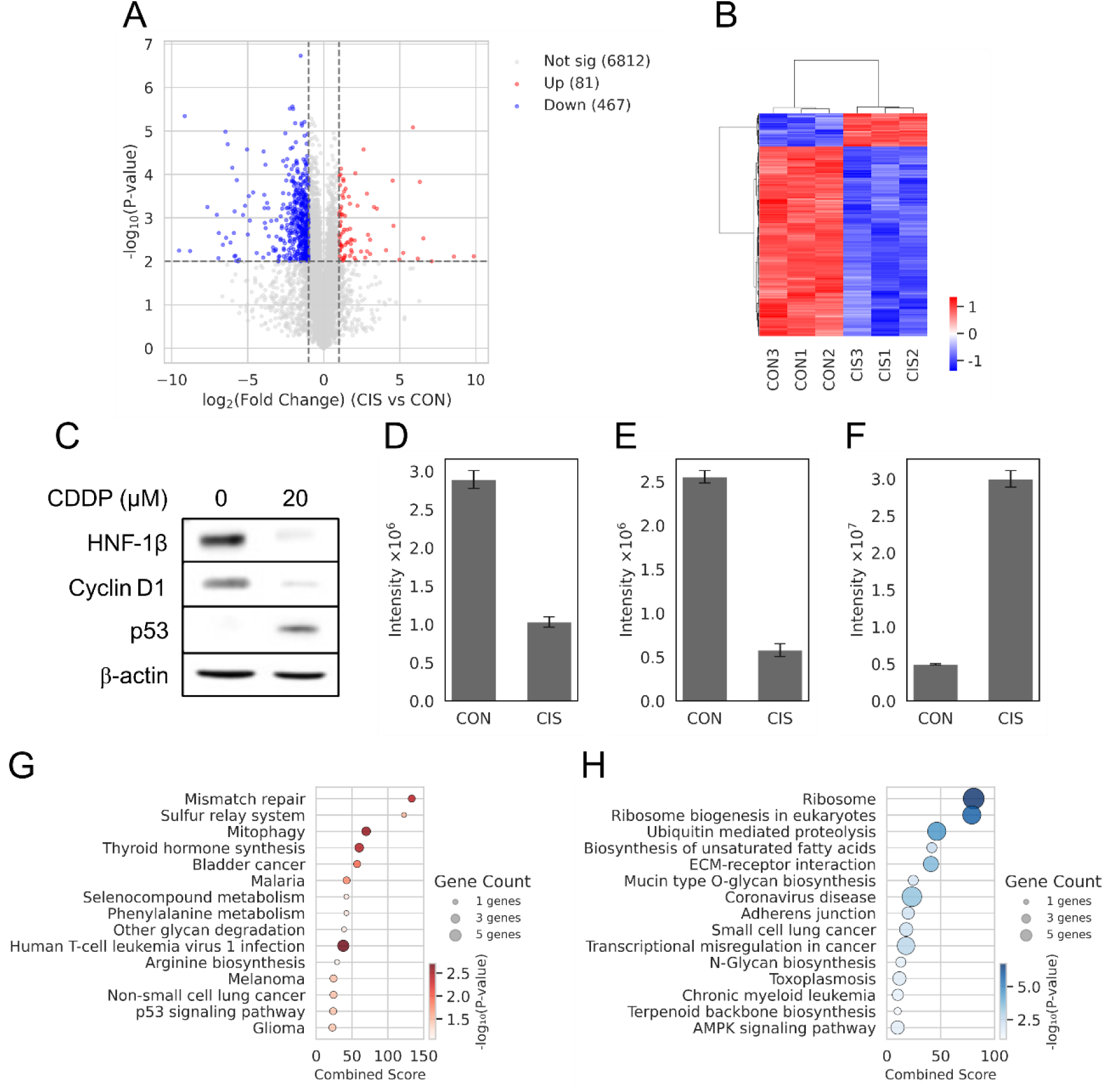
DIA-based quantitative comparison of OCCC cells with and without CDDP treatment. A. Volcano plot showing significantly upregulated, downregulated, and nonsignificant proteins between control (CON) and CDDP-treated (CIS) samples, using cut-offs of |log_2_(fold change)| > 1 and a *p* <0.01. B. Heatmap displaying proteins classified as upregulated or downregulated based on the volcano plot. C. Western blot validation of HNF-1β, Cyclin D1, and p53 in OCCC cells with and without CDDP treatment, using housekeeping β-actin as an expression control. D-F. MS intensities of triplicates for (D) HNF-1β, (E) Cyclin D1, and (F) p53 in OCCC cells with and without CDDP treatment. G–H. Enrichment analyses of (G) upregulated and (H) downregulated proteins showing the top 15 KEGG pathways, respectively.

## Discussion

IAA treatment induces nonspecific alkylation not only cysteine residues but also at other amino acids and at the N-terminus, generating multiple alkylated variants from the same peptide. These offsite modifications increase MS spectral complexity and diminish signal intensity (6, 7, 9). In contrast, DMSO functions as an oxidizing agent and promotes both thiol oxidation and disulfide exchange (12–14). Building on this property, we applied an oxidation reaction of DMSO coupled with the thiol-containing reagent 2-ME. As our previously reported, the 2-ME/DMSO workflow markedly suppressed these offsite modifications, directly addressing the fundamental limitation of IAA-based alkylation. Thereby, the workflow reduces MS spectral complexity intended for purified single proteins (9).

In this study, we applied this workflow to the proteomic samples, and 2-ME treatment markedly increased the number of identified peptides, with cysteine-modified peptides reaching approximately 1.6–1.9 times higher levels than those observed with IAA treatment (Figs. 1A and 2B). Previous report suggests that alkylation-induced increases in peptide hydrophobicity can enhance ionization efficiency in electrospray ionization MS (29, 30). While both IAA and 2-ME introduce alkyl modifications, the larger mixed disulfide adducts formed by 2-ME may result in greater increase in peptide hydrophobicity. Consistent with this notion, 2-ME-modified peptides in our dataset generally eluted later and produced stronger ion signals than their IAA counterparts (Supplementary Fig. S6). In addition to the signal-to-noise improvement, specific modification of cysteine residues led to improved quantitative reproducibility of cysteine-modified peptides (Supplemental Fig. S4A). Moreover, suppression of offsite modifications by 2-ME reduced MS spectral complexity and substantially increased the proportion of peptides quantified with CV ≤ 20% compared with IAA (Figs. 3D and Table 1).

Furthermore, when applied to a biological use case–cisplatin-treated OCCC cells–the 2-ME workflow maintained its high quantitative reproducibility, enabling the detection of pathway-level responses with confidence. The biological findings validated expected responses such as activation of p53 signaling and mismatch repair, as well as downregulation of ribosome-related proteins (31). Notably, these drug-responsive changes were detected in cysteine- and lysine-rich proteins such as CCND1, TP53, EXO1, as well as ribosomal proteins enriched in basic amino acids. Although the specific behavior of individual peptides was not directly examined in this study, this observation is consistent with the notion that suppression of offsite modifications and improved signal quality under the 2-ME/DMSO workflow may facilitate more reliable quantitative comparisons for proteins enriched in reactive amino acid residues. Together with western blotting and pathway enrichment results (Fig. 4), these findings support the utility of the 2-ME/DMSO workflow for accurately evaluating anticancer drug response. Importantly, approximately 90% of identified proteins were quantitatively analyzed with CV ≤ 20% under the 2-ME/DMSO workflow (Supplementary Fig. S7 and Supporting Table S1). These results indicate that the 2-ME/DMSO workflow provides a robust basis for practical quantitative proteomics applications.

## Conclusions

Our results demonstrate that 2-ME treatment substantially improves the reproducibility of peptide identification and quantification, especially for cysteine-modified peptides, across diverse proteomic workflows. These findings establish 2-ME adduction as a robust and practical strategy for high-precision and highly reproducible bottom-up proteomics. Moreover, the ability of 2-ME to suppress offsite modifications and reduce spectral complexity provides clear advantages for applications requiring accurate spectral interpretation, including peptide mapping or top-down proteomics. The simplicity of the workflow, together with its compatibility with diverse MS platforms, underscores its strong potential for broad adoption in proteomics research. Future work will be required to evaluate its performance in more complex applications, such as large-scale cohort studies and clinical proteomics, where simple and highly reproducible sample preparation is essential.

## Data Availability

The MS-based proteomics data including raw data files were deposited in the ProteomeXchange Consortium (http://proteomecentral.proteomexchange.org) via the jPOST partner repository (http://jpostdb.org)(32) as identifiers PXD071904 for ProteomeXchange and JPST004235 for jPOST.

## Supplementary data

This article contains supplemental data.

## Acknowledgments

This work was supported by Grants-in-Aid for Scientific Research from the Ministry of Education, Culture, Sports, Science and Technology, Japan (grant number 21K06036), and by Kitasato University (All Kitasato Project Study).

## Author contributions

A. S., Y. I., Toshihide. M., and M. I. investigation; A. S. formal analysis; A. S. and Takashi. M. conceptualization; A. S. methodology; A. S. and Takashi. M. writing–original draft; A. S., Y. K., and Takashi. M. writing–review and editing.

