## Supplementary material for "2-Mercaptoethanol/DMSO workflow enables highly reproducible quantitative proteomics": Supplemetary Information

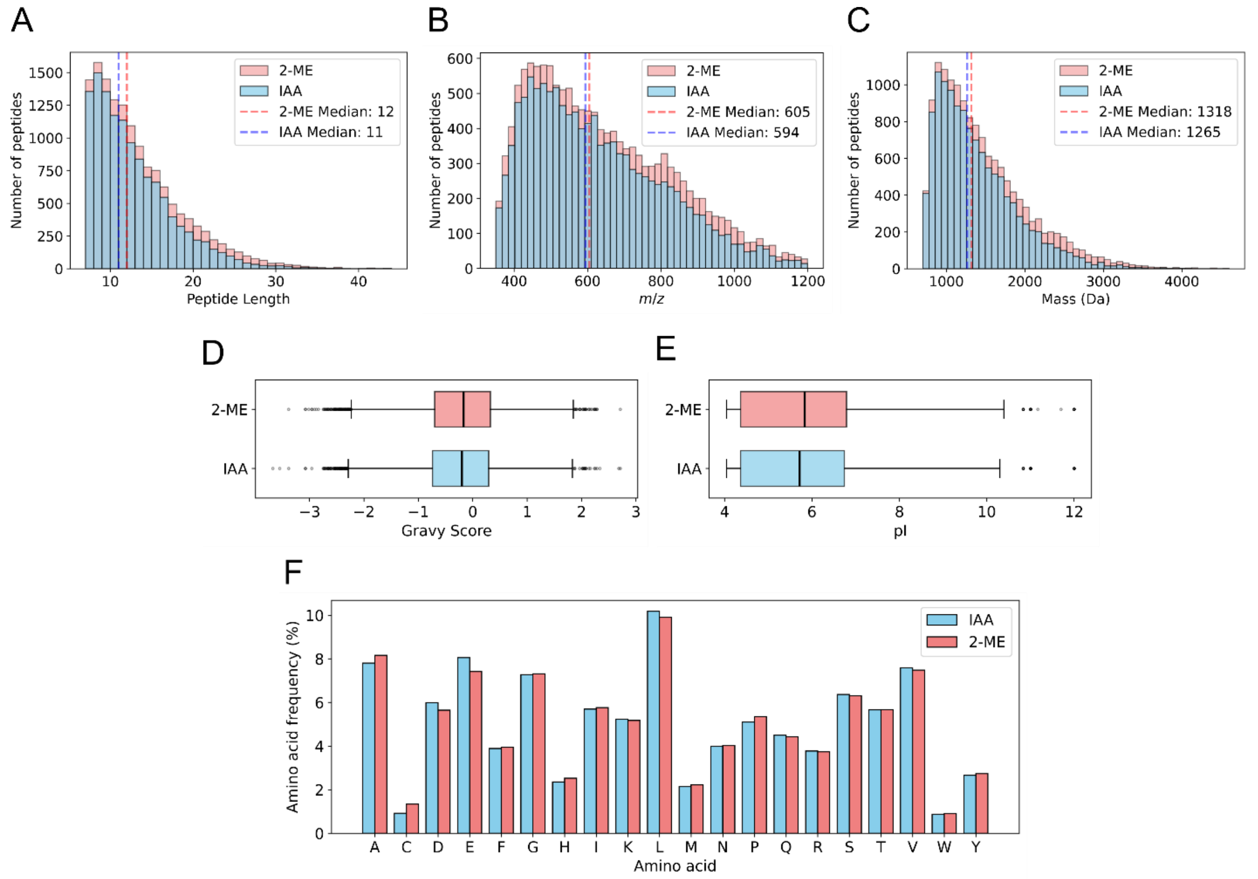


**Figure S1**. Comparison of physicochemical properties of identified peptides by IAA or 300 mM 2-ME. Panels A–F show the distributions of (A) peptide length, (B) *m/z*, and (C) molecular mass (Da), and the box plots of (D) GRAVY score and (E) pI, and (F) amino acid frequency profile, respectively. Peptides from IAA- and 2-ME-treated samples are shown in blue and red, respectively. Boxes indicate the interquartile range (IQR) between the 25th and 75th percentiles, respectively. Median value is depicted by black line. Minima and maxima within 1.5 times the IQR below the 25th percentile and above the 75th percentile are shown by whiskers. Outliers, signified by individual dots, fall outside the bounds.

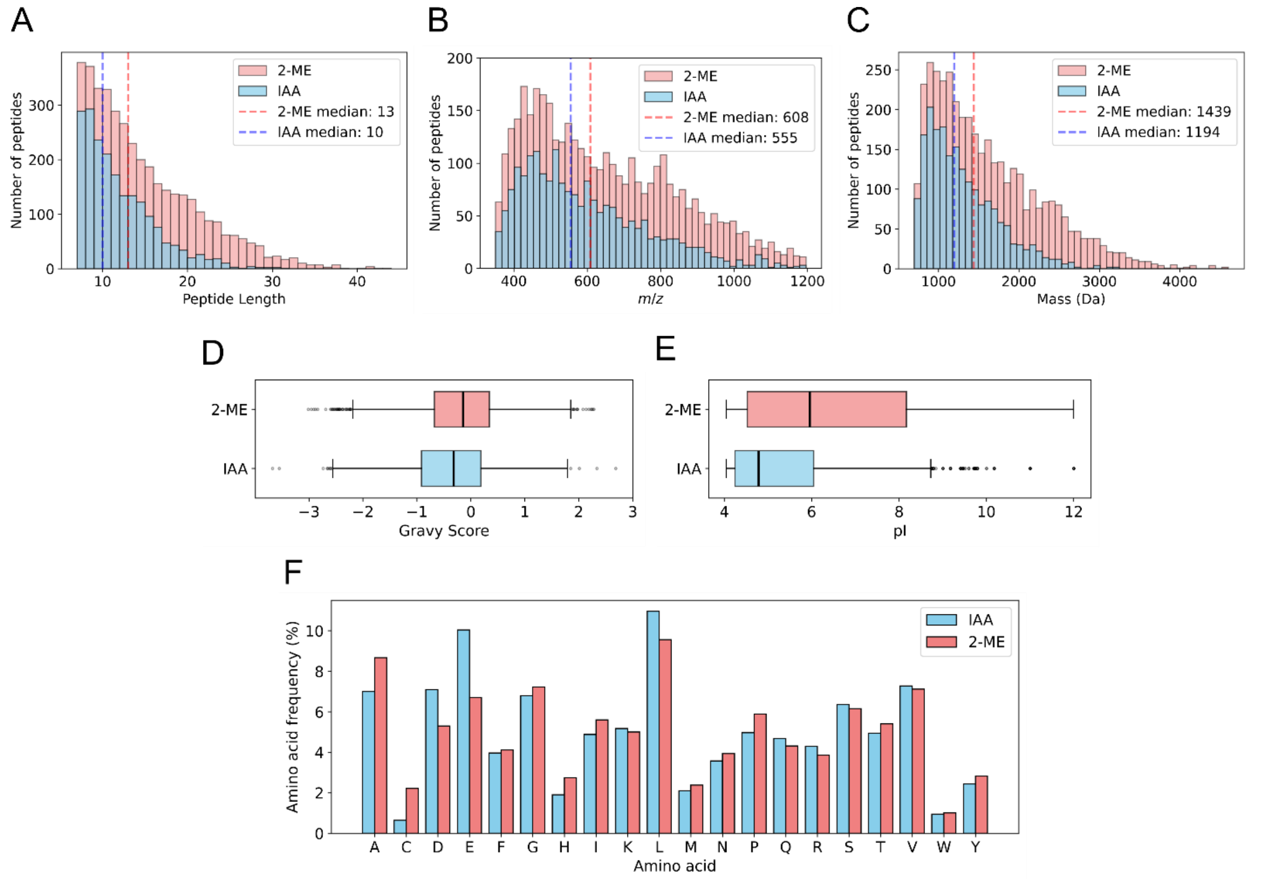


**Figure S2**. Comparison of physicochemical properties of peptides uniquely identified by IAA or 300 mM 2-ME. Panels A–F show the distributions of (A) peptide length, (B) *m/z*, and (C) molecular mass (Da), and the box plots of (D) GRAVY score and (E) pI, and (F) amino acid frequency profile, respectively. This figure is displayed in the same manner as Figure S1.

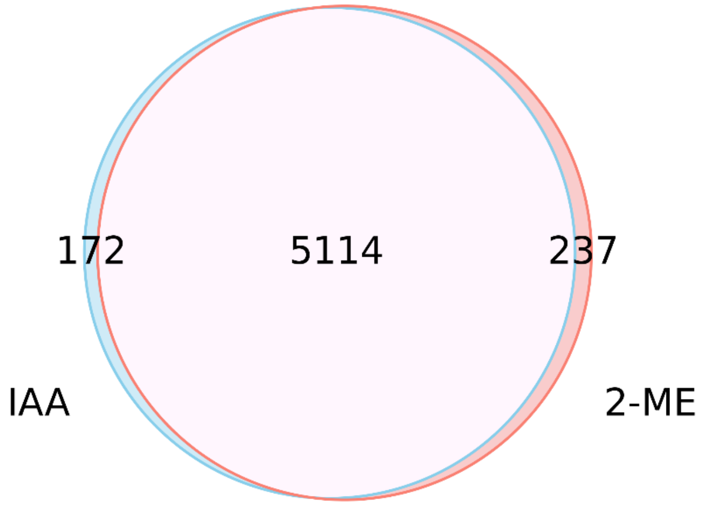


**Figure S3**. Ven diagram showing the overlap of proteins identified from IAA- or 300 mM 2-ME-treated samples. A total of 172 and 237 proteins were uniquely identified in the IAA- and 2-ME-treated samples, respectively.

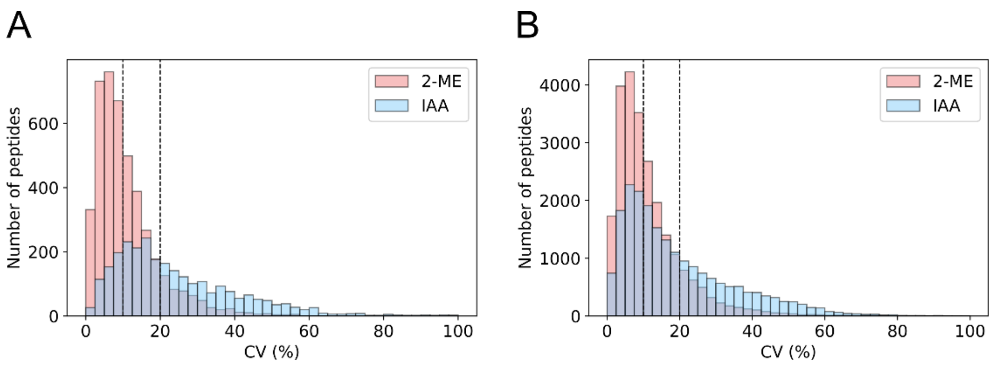


**Figure S4**. Distributions of CV values for peptides treated with IAA or 300 mM 2-ME. Panel A and B show cysteine-modified peptides and peptides without modified cysteine residues, respectively. Peptides from IAA- and 2-ME-treated samples are shown in blue and red, respectively. CV values at 10% and 20% depicted by dotted lines, respectively.

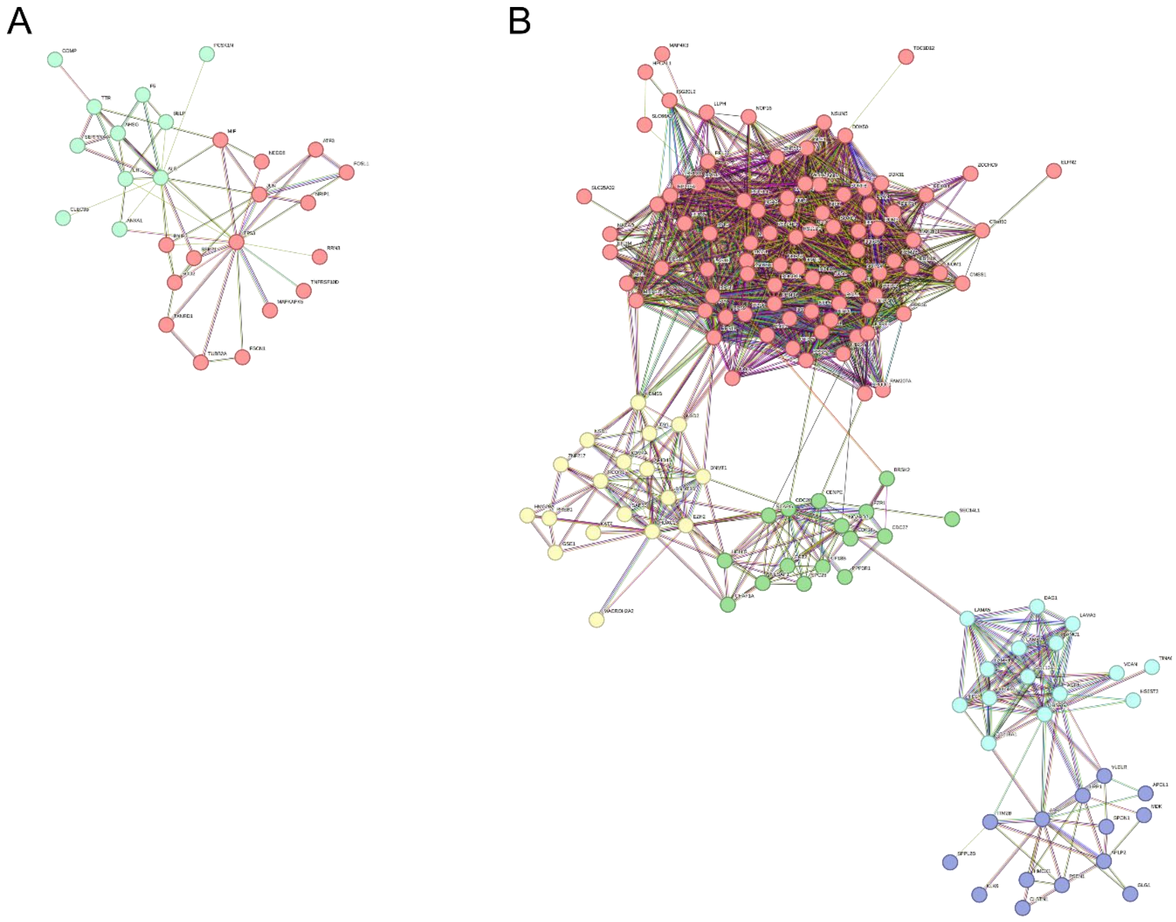


**Figure S5**. PPI networks generated by STRING analysis for differentially expressed proteins. Panels A and B show the networks for upregulated and downregulated proteins, respectively. Each color represents a different functional cluster of proteins within the network.

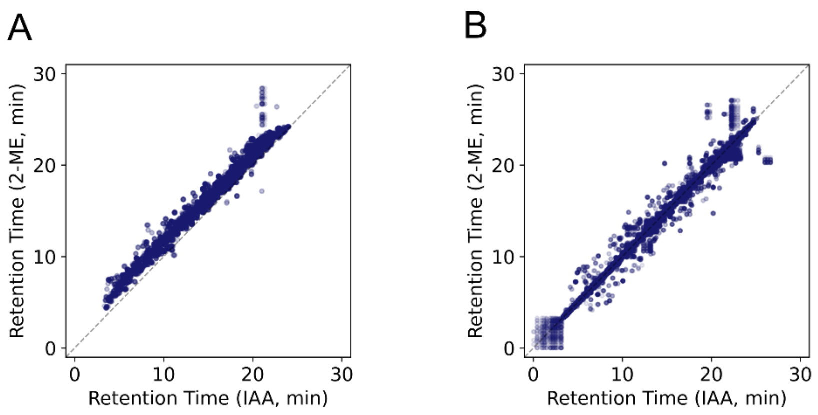


**Figure S6**. Comparison of peptide retention time between IAA and 300 mM 2-ME. Panel A and B show cysteine-modified peptides and peptides without modified cysteine residues, respectively.

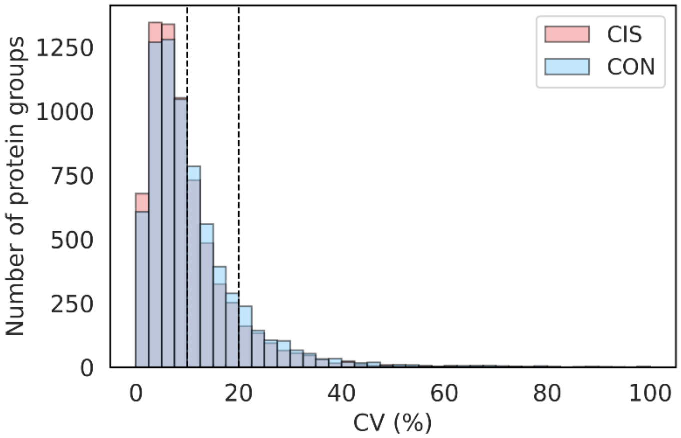


**Figure S7**. Distribution of CV values for quantified proteins comparing CON and CIS. Protein from CON and CIS is shown in blue and red, respectively. This figure is displayed in the same manner as Figure S4.

**Table S1**.CV of quantified proteins in OCCC comparing CON and CIS.

| | CON | CIS |
|---|---|---|
| CV ≤ 10% | 4,198 (59.0%) | 4,412 (64.0%) |
| CV ≤ 20% | 6,221 (87.4%) | 6,202 (90.0%) |
| CV > 20% | 893 (12.6%) | 690 (10.0%) |
